# Profiling extremophile bacterial communities recovered from a mining tailing against soil ecosystems through comparative genome-resolved metagenomics and evolutionary analysis

**DOI:** 10.1101/2024.08.28.610100

**Authors:** Moises A. Rojas, Gladis Serrano, Jorge Torres, Jaime Ortega, Gabriel Gálvez, Emilio Vilches, Valentina Parra, Angélica Reyes-Jara, Vinicius Maracaja-Coutinho, Lorena Pizarro, Mauricio Latorre, Alex Di Genova

## Abstract

Microbial communities inhabiting mining environments harbor a diverse array of bacteria with specialized metabolic capacities adapted to extreme conditions. Here, we utilized comparative genome-resolved metagenomics of a high-quality Illumina-sequenced sample from the Cauquenes copper tailing in central Chile. We investigate the metabolic roles and evolutionary behaviors of the resident microorganisms, focusing on capacities related to copper, iron, and sulfur metabolism. We recovered 44 medium and high-quality metagenome-assembled genomes (MAGs), primarily classified belonging to phylum *Actinobacteriota* (21), *Proteobacteria* (10), and *Acidobacteriota* (6). These MAGs were compared to the Global Soil MAGs project (SMAG catalog), which includes bacteria from conventional or natural ecosystems, to uncover specialized properties of mining bacteria. Notably, we discovered a new phylum, *Nitrospirota_A*, and provided insights into the unexplored taxonomic classifications at the lowest ranks such as genus and species. Functional potential analysis revealed that the mining community has enhanced molecular capabilities associated with sulfur and copper metabolism. Evolutionary analysis revealed that mining genes involved in targeted metabolism are under strong negative selection, indicating conservative evolutionary pressure within the mining environment. In particular, it was possible to identify a MAG from the genus *Acidithrix* with a global dN/dS ratio greater than 1, suggesting positive selection. Additionally, core proteins essential for bacterial survival, such as flagellar motors, cell cycle regulators, and biogenesis proteins, were also under positive selection. The latter points to the need for enhanced mobility in these microorganisms to locate resources efficiently. We demonstrate that copper mining communities are diverse and possess a significant metabolic repertoire under extreme conditions in sulfur and copper proteins. Those specialized genes appear to be in a conservative state rather than undergoing adaptive evolution. This study enhances our understanding of extremophile mining microbiomes, highlighting their high variability in classification, metabolic functions, evolution, and adaptation, which can be leveraged for further biotechnological applications.

## Introduction

Metagenomics aims to uncover the genetic information of complex microbial communities and their coexistence with other species interacting within a particular environment. Additionally, it enables the reconstruction of “metagenome-assembled genomes” (MAGs), *in-silico* genomes that cannot be obtained through the conventional isolation of microorganisms [1]. In recent years, metagenomics has revealed a valuable understanding of the complexity of heterogeneous microbial communities inhabiting extreme environments and their adaptations across various ecosystems characterized by low pH, wide range of temperatures, hypersalinity or high heavy metal concentrations [2][3][4][5]. Metagenomic analysis can predict taxonomic composition, metabolic-biogeochemical cycling functionalities, and selective pressure analysis of genetic evolution within these communities [6][7]. It is well-known that sulfur pathways dominate in acidic soils and hydrothermal environments [8][9], whereas nitrogen-carbon elements are prominent in agricultural soil microbiomes [10]. In particular, mining ecosystems, such as mining tailings also display an acidic behavior and are important resources as they are largest reservoirs of complex and unexplored microbial communities [11].

Some studies have explored microbial interactions using 16S rRNA sequencing of soil samples extracted from different copper tailing zones of the Cauquenes tailing [12]. The uncultivated microbial diversity of mining environments possesses unique capacities. For instance, the prokaryotic acidophilic *Acidithiobacillus thiooxidans* (a member of the genus *Acidithiobacillus*, a group of sulfur-oxidizing acidophilic bacteria) is able to tolerate acidic and heavy metal conditions under complex molecular pathways [13][14][15], making it a model mining microorganism in mining microbiology [16]. Over the years, other acidophiles, including those from genera *Leptospirillum*, *Acidimicrobium*, *Ferromicrobium* and *Sulfobacillus* have been described with similar behaviors and potential [17][18]. Moreover, metagenomic approaches of mining-derived MAGs have reported a consortium of microorganisms with relatively low taxonomic diversity that are involved in oxidative processes of iron and sulfur cycling, attributed to their unique genetic elements [19][20]. Those microorganisms hold potential for various applications, including the discovery of new genes, enzymes, or metabolic mechanisms [21]. They are recognized as prototypes for biotechnological applications in many countries, particularly for their efficient copper bioleaching capabilities, which are specially relevant to the Chilean mining industry [11][22][23]. Additionally, they are being considered for broader applications, such as the bioremediation of heavy metals and other water or soil pollutants [24][25][26]. Recently, extreme acidophiles such as those from mining environments have gained special attention for the generation of novel genetic engineering strategies [27].

Furthermore, the diversity of soil microbiome is not only complex, harboring valuable metabolic capacities and extensive genetic resources, but also largely unexplored at finer taxonomic levels (e.g., genus, species) [28][29][30][31], resulting in a significant portion of the bacterial composition still unidentified. It is key to deepen the understanding of microbial diversity within extreme environments through genome-resolved metagenomics, specifically focusing on a copper mining ecosystem as a model. In these environments, bacterial communities are often dominated by the phyla *Acidobacteriota*, *Actinobacteriota*, and *Proteobacteria*, as reported in mining and acidic soil studies [32][33]. In this context, the diversity of mining MAGs is expected to be considerably lower than in other or conventional soils, reflecting the extreme complexity and harsh conditions in which microorganisms inhabit [16]. This leads to a significant challenge, as only few studies have successfully reconstructed mining MAGs that meet even medium-quality standards [21]. Despite these challenges, the interest in metabolic pathways from metagenomic data has allowed the identification of key microorganisms and novel taxonomic groups within specific communities [34]. Nevertheless, substancial gaps persist in our understanding of the geochemical processes and novel taxa involved in these unique extreme environments. There is a growing interest in exploring how microbial communities adapt to acidic conditions of mining soils, particularly concerning the mechanisms they employ and their responses to such extreme environments. These aspects can be further elucidated through integrated multi-omics analyses [35].

In this study, we analyzed a metagenomic sample (labeled “S15”) collected from the Cauquenes copper tailing, one of the oldest (storing copper since 1936) and the largest copper waste deposit worldwide, covering 12.5 km^2^ in the O’Higgins region, central Chile. The Cauquenes deposit is characterized by high concentrations of metals (mainly copper, iron, zinc, minerals and salts) and an extremely acidic pH (∼3.5) [12]. Within this extreme scenario, we identified and characterized 44 medium to high-quality MAGs, profiling their taxonomic diversity and functional capacities in comparison to the recently reported Global Soil MAGs project (SMAG) [31], a comprehensive metagenomic catalog containing 32,931 soil bacterial MAGs from nine different ecosystems. We found the presence of a new phylum among the conventional ecosystems (*Nitrospirota_A*) and classified only 3 out of 44 as known species-level genome bins (or kSGBs) and 21 out of 44 as known genus-level genome bins (or kGGBs). This underscores the complexity and diversity of these microbial communities which are predominantly composed of previously unknown microorganisms. Moreover, the mining microbiome exhibited distinct metabolic capacities, particularly in copper and sulfur metabolism. Finally, an evolutionary analysis using dN/dS ratios revealed strong negative selection within intrinsic mining pathways, indicating it’s a conservative evolutionary state rather than one driven by adaptive pressure. Our findings enrich the current understanding of MAGs derived from mining microbiomes and suggest that, in opposition to previous reports, these mining microorganisms have high taxonomic diversity and unique metabolic capabilities that have been shaped by evolution.

## Results and discussion

### MAGs assembly from short-reads

The shotgun metagenomic sequencing of the S15 sample yielded 103,790,393 raw read pairs, with 99.23% passing the quality filters (see **Supplementary Table 1**). These filtered reads were subsequently processed for MAGs *de novo* assembly using the standard nf-core/mag v2.2.1 pipeline [36], available in Nextflow [37]. After metagenomic binning and refinement, we successfully generated 44 MAGs. Of these, 36 were grouped into three categories following the standards of Minimum Information about Metagenome-Assembled Genomes (MIMAG) [38]. It’s the quality control (QC) results, summarized in **Figure 1a**, classified 30 high-quality MAGs (≥90% completeness, ≤5% contamination), 5 as medium-quality (≥50% completeness, ≤10% contamination), and only one as low-quality (<50% completeness, ≤10% contamination). The remaining 8 MAGs included 3 classified as almost high-quality (completeness >90%, contamination between 5-5.2%), and 5 genome bins with contamination levels ranging from 10 to 25.18% (**Supplementary Table 2**).

**Figure 1.**
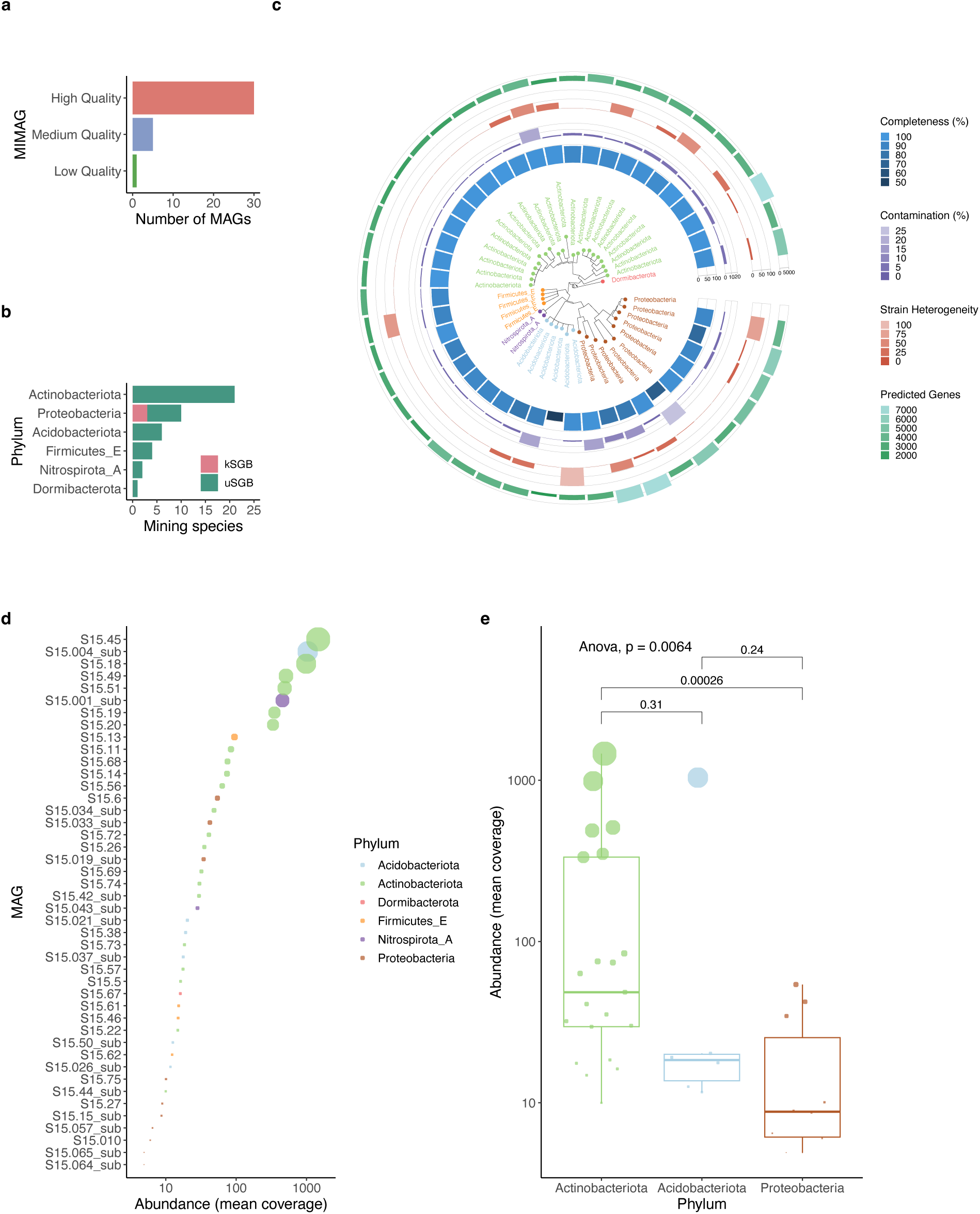
Quality metrics, taxonomic classification and abundances of the 44 MAGs obtained from the copper mining tailing. **a)** MIMAG classification of MAGs according to their genome completeness and contamination. **b)** Distribution of known/unknown species (kSGB/uSGB) at phylum level within the mining ecosystem. **c)** Phylogenetic tree showing the 44 MAG placements at phylum-level depicting individual assembly statistics (from inner to outer ring: completeness, contamination, strain heterogeneity, and number of predicted genes). **d)** Distribution of log10-transformed abundances (mean coverage of mapped reads) across MAGs, colored by phyla. **e)** Boxplot of abundances within the top three dominant phyla, sorted by median. ANOVA was applied to compare the means differences between the groups.

### Combined taxonomic composition of mining MAGs and the SMAG catalog

The taxonomic annotation of the 44 MAGs derived from the mining ecosystem was performed using the Genome Taxonomy Database Toolkit (GTDB-Tk) [39][40]. All MAGs were classified within the bacterial domain, with the phylum *Actinobacteriota* emerging as the most predominant clade (21 MAGs), followed by *Proteobacteria* (10) and *Acidobacteriota* (6), collectively encompassing 84% of the annotated phyla (**Figure 1b**). The phylogenetic tree, illustrating the phylum distribution and assembly statistics (completeness, contamination, strain heterogeneity) of the copper mining MAGs is depicted in **Figure 1c**. This dominance of specific phyla aligns with findings from conventional ecosystems (**Figure S1a**) and is consistent with the globally reported predominant phyla in soil microbiomes [29][31]. Moreover, all mining bins were assigned at the class level, 41 at the order level, 39 at the family level, and 21 out of 44 were classified as kGGBs. Only 3 MAGs were assigned as kSGBs: *Acinetobacter johnsonii*, *Pseudomonas_E hunanensis*, and *Sphingobium yanoikuyae* (see **Figure S1b**). All three belong to the phylum *Proteobacteria*, with two having medium quality and one having low quality. Interestingly, a new phylum was discovered in the metagenomic S15 sample: *Nitrospirota_A* (class *Leptospirillia*, order *Leptospirillales*, family *Leptospirillaceae*). This phylum, previously unexplored in mining-related microbiomes or the SMAG catalog of conventional ecosystems, includes two high-quality genomes (completeness >90% and contamination <2%), though it could not be annotated to either the genus or species level. Nevertheless, *Nitrospirota_A* has been reported as a polyphyletic group of *Nitrospirota* and comprises bacteria involved in nitrogen oxidative metabolism [41], and more recently in disproportionation reactions of thiosulfate and sulfur [42]. Abundances of mining MAGs are shown in **Figure 1d**, where the mean coverage of MAGs is highly dominated by *Actinobacteriota*. Interestingly, a high-quality MAG of the novel phylum *Nitrospirota_A* is also abundant (named “S15.001_sub”), regardless of its low representativity across known phyla. However, among the top three representative phyla (**Figure 1e**), *Proteobacteria* was found in low abundance, showing significant difference compared to *Actinobacteriota* (*p*-value = 0.00026).

Next, to compare the diversity of mining MAGs with those from other soils, we integrated them along with their associated metadata into the SMAG catalog, which is composed by 32,931 soil bacterial MAGs (*n* = 32,975 MAGs). A detailed analysis of the combined distribution across different environments revealed a consistent lack of classification at species-level genomes across all environments in different phyla (**Figure 2a**), which enforces the fact that knowledge of undiscovered microorganisms needs to be expanded, but still remains as a challenging matter especially those derived from extreme ecosystems [21][31]. In addition, the distribution of the known species ratio (the number of kSGBs divided by the number of MAG samples) across different ecosystems highlighted that artificial surfaces and grasslands are dominant, followed by mining (3 species in one sample). In contrast, bare lands, wetlands, forests, and shrublands exhibited the lowest ratios of annotated species (<0.6) (**Figure 2b**). Moreover, the kSGBs identified in the mining environment were also found in agricultural lands, artificial surfaces, bare lands, and grasslands, all within the phylum *Proteobacteria* (**Figure S1b**). **Figure 2c** illustrates all the kGGBs derived from mining that are shared across conventional ecosystems; for example the genus *Pseudomonas_E* was distributed across all environments except shrublands, whereas both *Sphingobium* and *Acinetobacter* were present in agricultural lands, artificial surfaces, bare lands, and grasslands; with the latter also found in wetlands. Other genera identified in the mining environment, such as *Ferrithrix* and *Acidithrix* were exclusive to mining, whereas *Rhodopila* and *Acidiferrobacter* were only shared with agricultural lands and grasslands, respectively.

**Figure 2.**
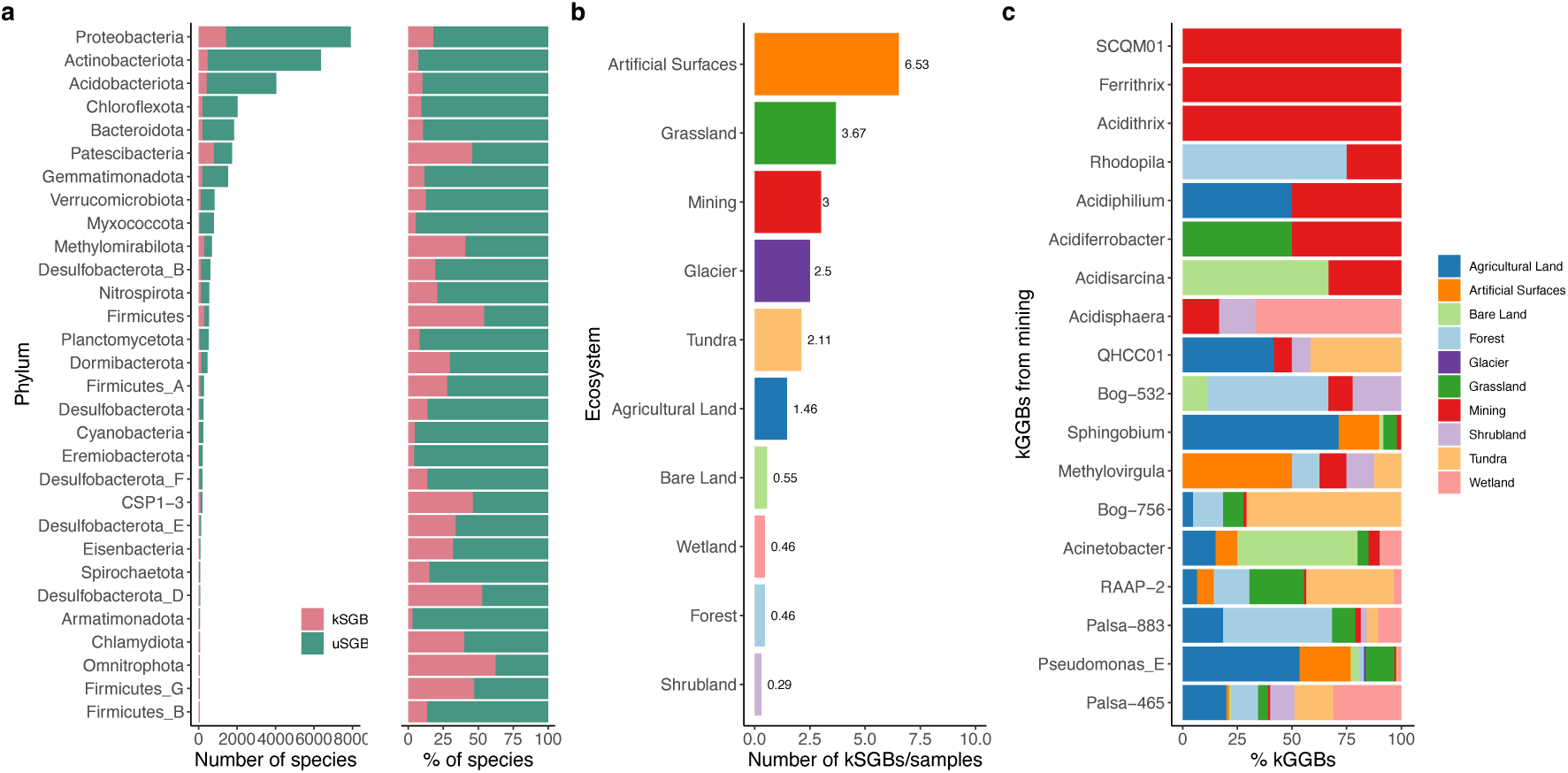
Combined taxonomic comparison between mining MAGs and the SMAG catalog. **a)** Distribution of known/unknown species (kSGB/uSGB) at phylum level (number and percentages of species), which is clearly dominated by uSGBs. **b)** Ecosystems ranked per ratio between the kSGBs and the number of metagenomic samples. **c)** Distribution of known genus (kGGBs) from mining that were also found across the conventional ecosystems, adjusted to 0-100%.

Although it is important to mention that the genus *Pseudomonas* has already been reported in extreme mining environments [12] with applications in various fields [43], this is the first time that *Pseudomonas_E* appeared in a metagenomic study linked to a mining ecosystem, whereas both genus *Sphingobium* and *Acinetobacter* have been previously identified in mining genomics [19][44]. Regarding the distribution of unknown species-level genome bins (or uSGBs) and their associated phyla, the most abundant across all ecosystems are governed as expected by *Proteobacteria*, *Actinobacteriota*, and *Acidobacteriota*, with the exception of artificial surfaces, where *Chloroflexota* and *Patescibacteria* emerged as dominant (**Figure S1c**). These dominant phyla are consistent with findings from other soil microbiomes using techniques such as 16S rRNA gene sequencing [45]. Overall, the taxonomic composition of mining metagenomics closely resembles those of conventional MAG soils at higher taxonomic levels of classification (e.g., phyla), but remains unique and less diverse at lower levels such as genus and species.

### Metabolic capacities of mining MAGs

For the purpose of examining the metabolic potential of mining MAGs and its comparison with the conventional ecosystems, we annotated the KEGG orthologs (KOs) compared to a representative subset of the SMAG catalog. First, we selected a set of 396 high-quality MAGs from conventional soils (sorted by completeness, grouped by ecosystem, non-duplicated based on their QC and classification parameters) to explore their functional landscape against the mining ecosystem (44 MAGs). A total of 7,044 unique KOs were comprehensively annotated across the 440 metagenomes using KOfamScan, with 4,258 KOs found exclusively in the mining environment. To further characterize these annotations within mining genomes, we assigned as selected pathways a total of 247 KOs that are directly involved in the intrinsic metabolism of copper, iron, and sulfur pathways (22, 68, and 157 marker genes, respectively). This analysis revealed several differences in the metabolic machinery of these biogeochemical cycles when comparing ecosystems.

Regarding copper metabolism, as exhibited in **Figure 3a**, the mining ecosystem ranks second after the glacier and along with artificial surfaces are the only ecosystems above the mean. Mining MAGs highlight the markers for copper resistance proteins (*CopB*, *CopC* and *CopD* genes) and *CsoR* family transcriptional regulator that controls copper oxidation and homeostasis (see **Figure S2a**). In the case of the iron proteins (**Figure 3b**), artificial surfaces are in the first position and proteins having transporter and binding functions are abundant across the other conventional ecosystems, but mining MAGs are placed last having a few outlier genes. Considering that iron mechanisms have mainly specialized in metal uptake under high competition scenarios, it is expected that in a mining environment, where there is high bioavailability of this micronutrient, there would be a lower need in terms of the quantity and diversity of proteins involved in iron acquisition. Interestingly, as depicted in **Figure 3c**, for sulfur proteins we found with statistical significance that mining MAGs are the top performers in this metabolism, highlighting the enzymes sulfide-quinone oxidoreductase (EC number: 1.8.5.4) and sulfurtransferase (EC: 4.4.1.37), along with the thiol-disulfide interchange protein (*DsbE* gene), and sulfur-carrier cysteine synthase and hydrolase proteins encoded by the *CysO* gene (EC: 2.5.1.113 and 3.13.1.6) that are associated to a mechanistic response of soil acidification [46][47]. Other sulfur transferases and regulatory proteins also dominate this intrinsic metabolism, as highlighted in the bubble plot in **Figure S2a**, underscoring the crucial role of these genes in the mining ecosystem compared to conventional ecosystems.

**Figure 3.**
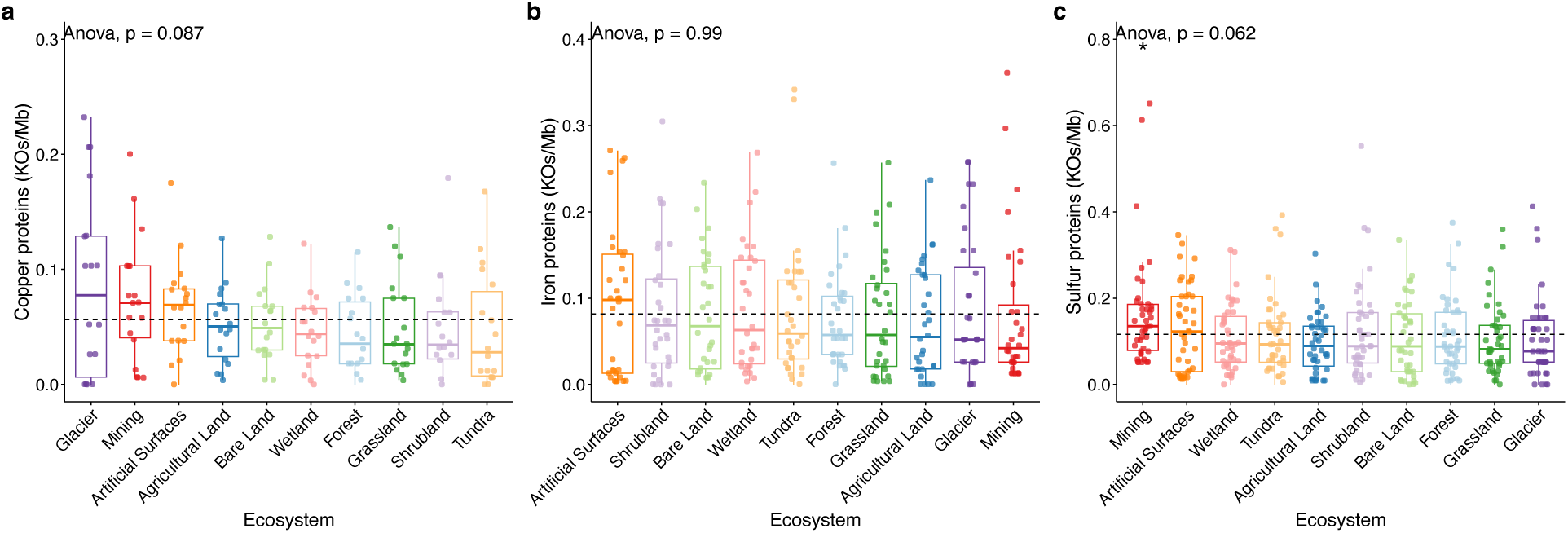
Functional metabolic capacities of mining MAGs compared to conventional ecosystems present in the SMAG catalog. The description of selected geochemical cycles and metabolic pathways potential was assessed using KOfamScan annotating the resulting KEGG orthology groups (KOs). Boxplots showing the KO definitions at selected **a)** copper, **b)** iron, and **c)** sulfur proteins sorted by ecosystem. Each dot represents a KO annotation where the counts were normalized using the length for all MAGs per ecosystem (KOs/Mb, y-axis). The *p*-value represents the statistical difference between ecosystems calculated with an ANOVA (* = 0.05 significance). In all plots, the ecosystems (x-axis) are sorted by median in decreasing order.

The selected KOs and their associated metabolic pathways were summarized in a heatmap (**Figure S2b**), where the 247 annotations were filtered by median (0.097), resulting in 124 genes. Upon examining the normalized abundances by z-score, no remarkable specialization was noticed in the mining MAGs for the iron proteins, which are primarily involved in transport (15/34 KOs). Moreover, for sulfur proteins (74 KOs), other ecosystems, such as forests, grasslands, and wetlands exhibited distinct activity profiles based on their abundances. Their metabolism is more distributed, comprising reduction, oxidation and transferase capacities (22, 15, and 9 genes, respectively). Finally, the copper proteins (16 KOs) are governed by metal resistance genes and copper carriers (4 and 3, respectively). In general, the availability of nutrients like nitrogen, iron, copper, or zinc is a component of plant enzymes for their uptake and transport; therefore are considered as key nutrients of a wide diversity of benign soils [23][48][49][50], fact that is clearly noticed in the heatmap. It is worth mentioning that the capacities of mining MAGs are enriched in uSGBs, resulting in significant intrinsic metal oxidative-reductive reactions (iron and sulfur), as well as for copper resistance, a fact that emphasizes the relevance of mining genomes in this pathway. Although annotations show slight differences in certain mining MAGs compared to other soil ecosystems, interpreting and comparing them is visually complex.

In addition, an specific metabolic-biogeochemical analysis was performed in the extremophile MAGs from mining using MEBS (Multigenomic Entropy-Based Score) [51], a tool that integrates curated pathways for known cycles and protein families from the Pfam database [52] to understand the diversity of certain microbial taxa inferred by using an entropy score for likelihood. As exhibited in **Figure S2c**, we found a similar response across the mining genomes where the most likely reactions involved are sulfur oxidation (*DsrEFH*, *FccB*, *Sox* genes), reduction (*sreABC*, *QmoABC*, *DsrABC*), disproportionation (thiosulfate), assimilation (*CoB*/*CoM*), and also nitrate reduction (denitrification, assimilatory). These findings are in accordance with previous reports on the sulfur pathway detected in acidic microbiomes [33][53][46][47]. Acidic environments like copper mining tailings provide a great habitat for sulfur-oxidizing microorganisms that require, for instance, the enzyme molybdopterin sulfur reductase (*sreABC* gene), which is needed for elemental sulfur (S^0^) reduction to H_2_S for energy production [54]. Besides, the abundance of this compound is consistent with the presence of genes encoding enzymes involved in the oxidation of a variety of reduced sulfur compounds (H_2_S, S^0^). This gene was particularly found in the majority of mining MAGs with medium to high likelihood. On the contrary, marker genes belonging to nitrogen metabolism are annotated with low abundances, as they are expected in other ecosystems like forests or cultivable lands [55].

### Evolutionary analysis of mining MAGs

In order to assess evolutionary pressures acting on the 44 assembled mining MAGs, we used the dndscv package to estimate the ratios between the number of nonsynonymous and synonymous substitutions (dN/dS) per gene and globally per MAG. We identified a total of 146,932 high-quality SNPs, estimating the global dN/dS ratios for a total of 13,601 genes. The global dN/dS ratios for the 44 mining MAGs, where each per-gene ratio was normalized by percent of contamination (**Figure 4a**), indicate a strong negative selection operating on this ecosystem. Moreover, **Figure 4b** shows the average estimation of dN/dS grouped at the top three dominant phyla, which resulted in phylum *Acidobacteriota* exhibiting significant average difference compared to both *Proteobacteria* and *Actinobacteriota* (*p*-value = 0.023 and 0.0097, respectively). Interestingly, we identified a single outlier MAG (named “S15.49”) with a global dN/dS of 1.27 that was classified within the genus *Acidithrix* –a particular genus that we found only in the mining ecosystem (as previously shown in **Figure 2c**)– and was assembled at high-quality (98.72% completeness, 1.28% contamination; as represented in **Figure S3a**), therefore being the one involved in adaptive evolution among the mining MAGs; in contrast with the MAG corresponding to *Acidobacteriaceae* family that has exhibited the lowest dN/dS ratio of 0.17 implying strong negative selection associated to their gene pool. In particular, *Acidithrix* is a predominant genus in mining scenarios inhabited by acidophilic bacteria [56][57][58]. Its absence in the other analyzed environments indicates that this genus has primarily specialized in inhabiting mining settings. This is consistent with its greater genetic plasticity, which allows it to adapt to this environment more effectively compared to other species found. This particular MAG is highly represented within the sample as it ranks fourth among the abundances distribution of mapped reads across MAGs (**Figure 1d**).

**Figure 4.**
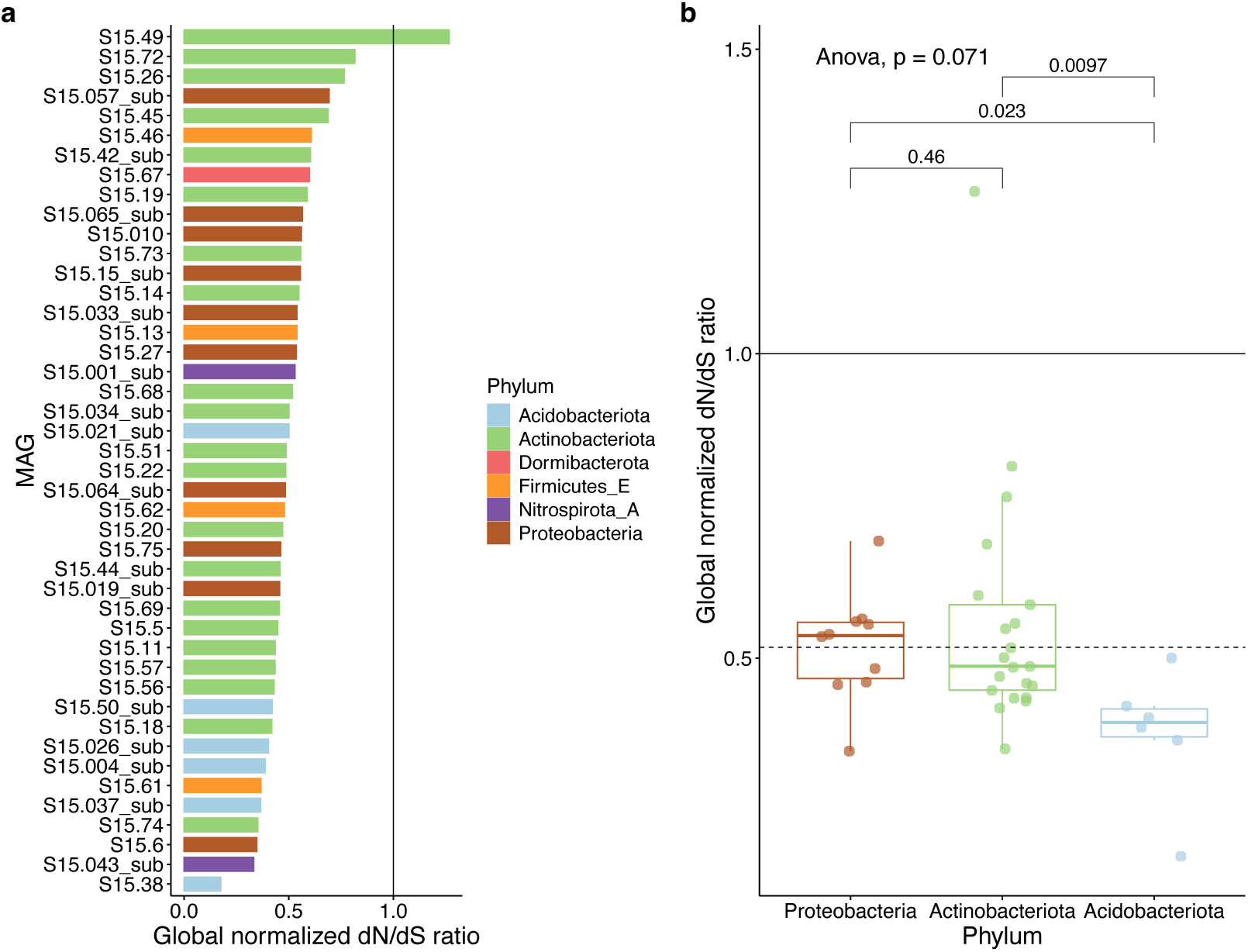
Global dN/dS ratios of mining MAGs and its distribution across phyla. **a)** Barplot for global estimation of dN/dS ratios (normalized by MAG contamination) accumulated in the 44 MAG bins, sorted by descending strength and colored by phyla. **b)** Boxplot of global estimation of dN/dS ratios within the top three representative phyla, where each dot represents a MAG. Ratios were normalized by MAG contamination, then grouped and sorted by median. ANOVA was applied to compare the means differences between the groups.

At the gene level, a total of 12,067 genes have a dN/dS ratio <1 (88.7% of total) associated with purifying or negative selection (NS). Based on our results, we found 84 genes under NS that are involved in the selected pathways that were previously called by the KOfamScan annotation. In particular, the number of annotated CDS is ranked as follows: iron, copper, and sulfur proteins (30, 28, and 26 genes, respectively). Those genes have purifying selection at appreciable rates (overall mean of 0.197), as depicted in **Figure 5a**, where iron exhibits the most conservative metabolism with significance over sulfur (*p*-value = 0.008). The mean dN/dS values for each pathway were 0.273 for sulfur, 0.210 for copper, and 0.120 for iron. In addition, this list of 84 genes was grouped by gene products and ranked by means of dN/dS ratios. Thus, **Figure 5b** shows 39 unique gene products, where the top three proteins are involved in the metabolism of sulfur (the enzyme sulfurtransferase, *CysO* and *SufE* genes).

**Figure 5.**
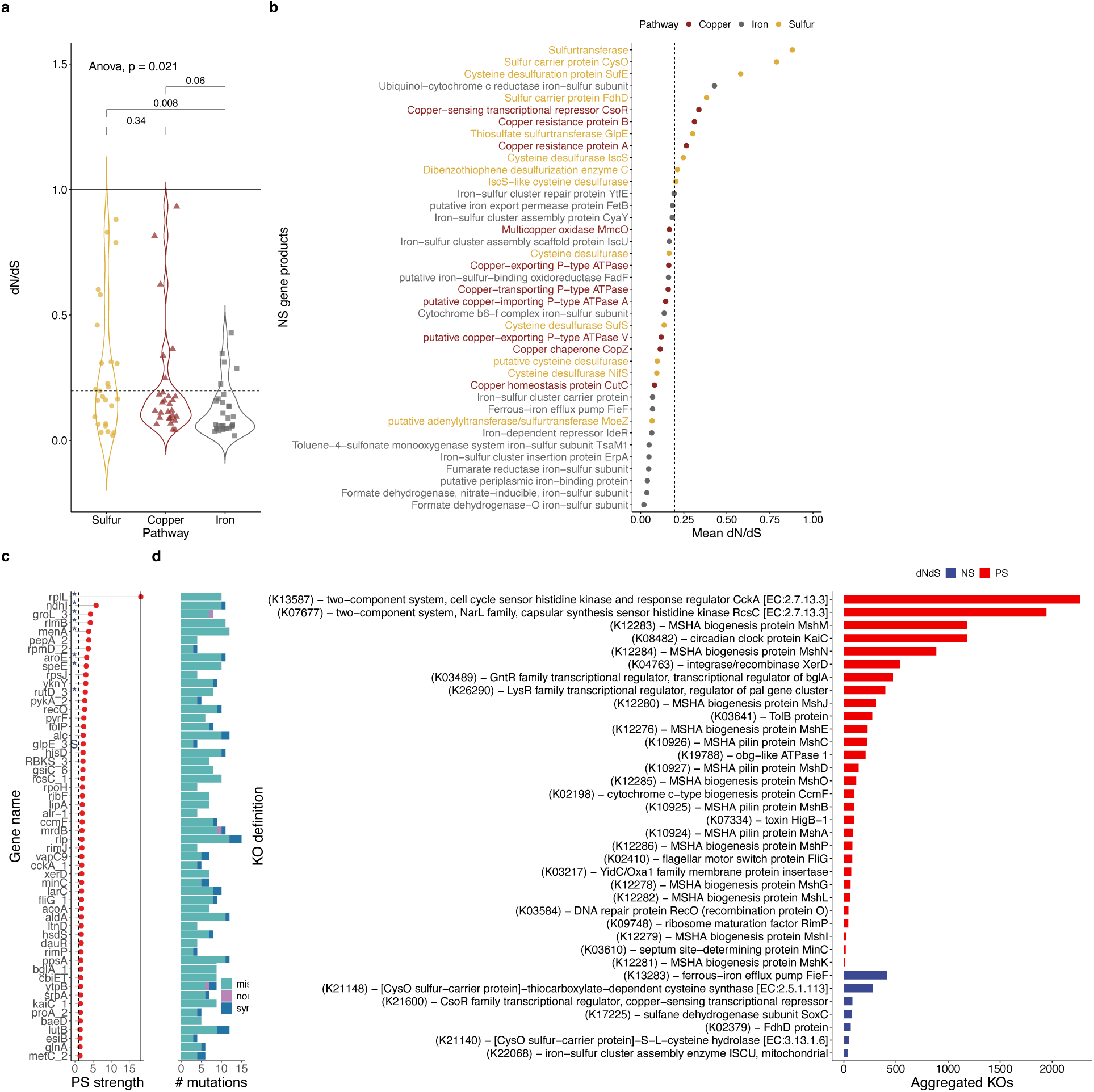
Evolutionary analysis on mining MAGs, summarizing the genes under positive (PS) and negative selection (NS). **a)** Violin plot for genes under NS focused on intrinsic mining pathways, ordered by decreasing dN/dS ratios (the dashed line represents the overall mean). An ANOVA was applied to differentiate between groups. **b)** Dotplot of the annotated gene products for selected NS genes, colored by pathway and sorted by selection strength. Each dot depicts the mean dN/dS among unique products (the dashed line represents the overall mean). **c)** Ranking of genes under PS and number of mutations per type, discarding hypothetical proteins and genes holding below 4 SNPs. Only for visualization purposes, a dN/dS ratio cutoff was set to 1.4. (legend: * = genes reaching FDR significance; S = sulfur protein; mis = missense mutations; non = nonsense; syn = synonymous). **d)** List of KEGG KOs and their definitions involving genes under both PS and NS (*n* = 36), reduced to unique KOs containing at least 10 aggregated genes.

Furthermore, a total of 1,534 genes have a dN/dS ratio >1 (11.3% of total) and are under positive selection (PS), widely exceeded by genes under NS. In detail, 266 of them have a false discovery rate of 5%, 1,370 (89.3%) were annotated as hypothetical proteins (HPs) and only 93 genes have at least 4 SNPs; those were used for further analysis (**Figure 5c**). It can be noticed that only 8 genes reached significance (*p*-value <0.05) where the *rplL* gene, which encodes the ribosomal subunit protein bL12, has the higher selection strength (dN/dS = 18.21). Sequence variations within rRNA genes can modulate ribosome function, influencing gene expression and cell physiology. This can be particularly advantageous for bacteria in extreme environments, where specific rRNA variants may provide a survival benefit [59]. In the case of the mining environment, based on the results obtained, it appears that mining species could adapt to extreme conditions through changes in protein levels while maintaining the same functions. In this context, only one of those genes was marked as related to intrinsic mining processes: the *glpE* gene, encoding the thiosulfate sulfurtransferase enzyme which participates in sulfur transfer reactions [60], that was found in a high-quality MAG assigned to phylum *Proteobacteria*. At species level, the total number of PS genes is summarized in **Figure S3b**.

The above results can also be correlated with the annotation regarding KEGG KOs, as depicted in **Figure 5d**, in which the PS and NS genes are collected regarding their aggregated number of KOs. To mention, proteins linked to iron and sulfur pathways are overrepresented by genes under NS (e.g., sulfur carriers: *CysO* and *FdhD* genes; iron efflux pump: *FieF* gene which mediates stress response to high concentrations of iron and zinc [61]); whereas the genes under PS dominate the core protein functions for bacterial survival such as flagellar motors, cell cycle regulators and biogenesis proteins, highlighting several KOs of MSHA-type motor proteins [62] suggesting the underlying mechanisms in which bacteria are evolving in the mining ecosystem. In mining environments, flagella play a crucial role in the initial stages of biofilm formation by allowing bacteria to move to a suitable surface and establish an organized structure [63], thus avoiding local high concentrations of metals.

Overall, the evolutionary analysis implies that genes under negative pressure of selection constitute an essential factor for the mining species due to their involvement in intrinsic metabolic pathways, indicating that those capacities are maintained in a conservative state rather than evolving. This phenomenon has been lightly studied in microbial communities across different ecosystems, where the remark is that the evolutionary rate in environmental metagenomes from extreme ecosystems is faster than in conventional environments [64]. Yet, there remains a surprisingly low number of studies specifically focused on mining ecosystems. Our findings suggest that the conservative evolutionary pressure shaping these bacteria may challenge the proposals for biotechnological applications aimed at improving or genetically enhancing specialized mining-derived processes, such as biomining or bioleaching for industrial production, due to the genes under PS are involved in bacterial functions mainly related to survival processes.

### Conclusions

We assembled and annotated 44 medium and high-quality metagenome-assembled genomes (MAGs) from a metagenomic sample of the Cauquenes copper tailing (S15). Taxonomically, we observed a prevalence of bacterial species belonging to the phylum *Actinobacteriota*, *Proteobacteria*, and *Acidobacteriota*, which were also found to be dominant in the SMAG catalog containing bacterial MAGs from nine different conventional soils. Additionally, we identified a new phylum, *Nitrospirota_A*. At the species level, only three out of the 44 MAGs were annotated: *Acinetobacter johnsonii, Pseudomonas hunanensis, and Sphingobium yanoikuyae*, and are shared with agricultural land, artificial surfaces, bare land, and grassland.

The metabolic machinery of the mining MAGs underscores the importance of copper and sulfur pathways proteins, highlighted by key genes, such as those involved in copper resistance and carrier/transporter proteins specific to acidic environments like copper mining tailings. In contrast, iron metabolism showed no relevant activity on mining compared to the other ecosystems. The evolutionary analysis using dN/dS allowed us to investigate genes under positive selection (PS) and negative selection (NS). Notably, only one gene related to the metabolism of sulfur was identified to be under PS (*glpE*), indicating a clear trend toward a conservative evolution in the intrinsic mining pathways. We also identified a high-quality MAG belonging to the genus *Acidithrix*, uniquely established within the mining ecosystem, which is evolving under a global positive selection.

We demonstrated that the evolution of the mining microbiome is skewed towards genes under NS across sulfur, copper, and iron proteins, with the iron metabolism showing a dominant conservative impact over dN/dS ratios. However, motor and biogenesis proteins are primarily governed by PS genes, suggesting enhanced capabilities of mining bacteria to locate resources. Overall, our study, leveraging genome-resolved comparative metagenomics, reveals that mining microorganisms are highly specialized in intrinsic pathways, paving the way for targeted biotechnological applications. The presence of unclassified species as novel extremophiles underscores the high, unexplored biodiversity within these ecosystems, which are under evolutionary pressure to preserve their specialized capacities.

## Methods

### Sample collection and DNA extraction

Three replicated soil samples (100 g each) were collected in July 2022 (Chilean winter) under sterile conditions at 5 to 10 cm of the soil surface in a particular zone of the Cauquenes copper tailing belonging to the intermediate site and labeled as S15 sample (pH ∼3.5, high copper concentration). The replicates were homogenized in situ before being stored on dry ice for further microbial analyses. DNA was extracted from the collected samples (each of the three soil samples) using the Zymo kit Quick-DNA Fecal/Soil Microbe MiniPrep^TM^, combining the manufacturer’s instructions and pre-treatment of the samples before following the kit protocol. 20 grams of soil were resuspended in 20 ml PBS buffer. Then, the sample was homogenized using FastPrep-24 equipment (MP Biomedicals Cat. Number 6004500) in a cycle of 4 m/s for 20 seconds. The mixture was centrifuged (120 × g for 5 min) at room temperature and the supernatant fluid was transferred to a clean tube, and centrifuged (7,100 × g for 10 min) at room temperature, then the supernatant was discarded and the pellet was resuspended in 1 ml of PBS buffer. After, the samples were vortexed for 10 seconds, and the mixture was transferred into a DNA Fecal/Soil Microbe lysis tube to continue the kit protocol. Concentrations of DNA were evaluated with a Qubit 4 fluorometer. Microbial DNA was amplified using a bacteria-specific primer set, 28 F (5’-GA GTT TGA TCM TGG CTC AG-3’) and 519 R (5’-GWA TTA CCG CGG CKG CTG-3’), flanking variable regions V1-V3 of the 16S rRNA gene [65], with barcode on the forward primer. Amplification protocol was performed using the promega GoTaq® G2 Flexi DNA Polymerase Mix, under the following conditions: initial denaturation at 94 °C for 3 min, followed by 28 cycles, each of them configured at 94 °C for 30 s, 53 °C for 40 s and 72 °C for 1 min, with a final elongation step at 72 °C during 5 min. After amplification, PCR products were checked in 2% agarose gels and finally stored at −20 °C.

### Metagenome sequencing

The extracted DNA from the S15 sample was later used for shotgun metagenomic sequencing to evaluate the microbial community inhabiting the copper mining ecosystem. According to the manufacturer’s instructions, metagenomic libraries were constructed using a TruSeq^TM^ DNA PCR-free Sample Prep Kit (Illumina, USA) according to the manufacturer’s instructions. A total of 103,790,393 read pairs of raw metagenomic reads were generated with a HiSeq^TM^ 2000 Illumina system (2 x 150 bp) (see **Supplementary Table 1**).

### Metagenomic assembly, binning, quality assessment, and classification

Raw reads were assembled, binned and annotated using the nf-core/mag v2.2.1 pipeline [36] available for the workflow management system Nextflow [37], on a 1,000 cores local cluster with 200TB storage and 7TB of RAM. First, fastp v0.23.2 was used to perform the short reads trimming to remove adapters and low quality reads [66]. Quality control of the trimmed reads was assessed with FastQC (v0.11.9). Co-assembly of the trimmed reads was performed using SPAdes v3.15.3 [67][68], binned with MetaBAT2 v2.15 [69] and MaxBin2 v2.2.7 [70], then pooled and refined employing DASTool v1.1.4 [71] setting a min contig size of 1,500. This computational methodology allowed us to retrieve a total of 188 genome bins reconstructed from the metagenomic sample (S15), but after refinement only 44 genomes were subsequently considered for downstream analysis. Additionally, we perform a comprehensive evaluation with the metashot/prok-quality pipeline running also in Nextflow [72], to estimate the genome quality of the refined bins by CheckM v1.1.2 [73]. The global average GC content is 62.14, N50 is 41,379 bp, and the number of contigs is 19,346. Gene prediction was carried out with Prodigal v2.6.3 [74], and the prediction of CDS and functional annotation of mining MAGs was performed with Bakta v1.8.1 running the full database [75]. Finally, the taxonomy classification was conducted with the Genome Taxonomy Database Toolkit (GTDB-Tk v1.5.0, database version r202) [39][40], resulting in a phylogenetic tree with MAG placements inferred using a set of 120 bacterial single-copy genes (no archaeal marker genes were found). The reference tree was built using the R package ggtree v3.10.1 [76].

### Functional annotation of MAGs and statistical analyses

KEGG orthologs (KOs) were predicted using the KOfamScan annotation tool [77] with default options. The protein sequences of the 44 mining MAGs and a representative subset of the SMAG catalog comprising 396 bins from the conventional ecosystems were queried against the HMM models from KEGG database (release 108.1, 2023). Then, the raw results were merged in a matrix with MAGs in columns and KOs in rows, which was further processed to normalize the countings divided by the total genome size for each ecosystem in order to build the boxplots for targeted pathways (Cu, Fe, and S) with the help of the R package ggpubr v0.6.0. This matrix was also used for the bubble plot with relevant genes from the mining ecosystem and for the annotated heatmap built with the R package ComplexHeatmap v2.18.0 [78]. Moreover, to evaluate if the means are equal between ecosystems, we include a global analysis of variance (ANOVA) in each boxplot. Next, we also evaluated the metabolic capacities of the mining MAGs with MEBS (Multigenomic Entropy-Based Score) v1.2 [51], a tool that integrates curated pathways for known cycles (naming C, Fe, N, O, S) and protein families from the Pfam database [52], aimed to understand the intra-diversity within the mining metagenomes by using an entropy score for likelihood. The resulting dataset with abundances was used to generate the heatmap relating geochemical cycles across the 44 MAGs.

### SNP calling

To perform variant calling in the metagenomic mining sample (S15), first the reads were mapped to the previously assembled bins (in FASTA format) using the aligner bwa-mem2 v2.2.1 [79], then the resulting alignment was converted from SAM format to a BAM file with samtools v1.16.1 [80]. This BAM file was processed using elPrep tool [81] in the split/filter/merge mode (sfm) to remove duplicates and then sorted by coordinate order (samtools sort command). The sorted BAM file and the reference genome were used as input for BCFtools to call variants [82], based on its mpileup algorithm output (default specifications) and the multi-allelic calling model in the call subcommand (-m option and ploidy option set to 1), directing all the variants to a VCF file. After, in order to evaluate the impact of the detected SNPs, we used the SnpEff v5.2a program [83] which needs two steps: first, to manually build a database for the MAGs using both the assembled bins and the information on where the genes are in the genome (a single-merged GFF file from the CDS annotation); and second, to perform the genomic annotations using this custom database and the VCF file with the variants (bcftools output). Finally, synonymous and non-synonymous mutations were analyzed using the R package dndscv version 0.0.1.0 [84], where we defined genes under positive selection (PS) as those having a dN/dS ratio >1, and genes under negative selection (NS) with a ratio below 1. The false discovery rate (FDR) was adjusted at a *p*-value below 0.05. Circos plot visualization was created using the package circlize v0.4.16 available for R [85].

## Code availability

The main datasets and code used in the analyses along with the figures generated for this work are available on GitHub (https://github.com/digenoma-lab/biomining_metagenomes).

## Author contributions

M.A.R. contributed to the design, data collection and processing, analysis and interpretation, literature review, writing, and critical review of the manuscript. G.S., J.T., J.O., and G.G. were involved in data collection and processing, and materials. E.V., V.P., A.R.-J., V.M.-C., and L.P. contributed to funding acquisition and critically reviewed the manuscript. L.P was also involved in data collection and processing. M.L. was responsible for the conception, supervision, funding acquisition, materials, data collection and processing, writing and critical review of the manuscript. A.D.G. contributed to the conception, design, supervision, funding acquisition, analysis and interpretation, literature review, writing, and critical review of the manuscript. All authors have read and agreed to the published version of the manuscript.

## Acknowledgments

This work was supported by Proyecto Anillo Regular ANID ACT210004; Center for Mathematical Modeling; Apoyo a Centros de Excelencia ACE210010; ANID Millennium Science Initiative Program ICN2021_044; Fondo Basal FB210005; ANID FONDECYT 1190742, 1221029, 1230195, 1211731, and 1230194; Fondo Interdisciplinario y Proyecto Núcleo UOH; Proyecto Postdoctoral ANID 3220080; supercomputing infrastructure of the NLHPC (ECM-02); the supercomputing infrastructure of the High-Performance Computing UOH laboratory (FIC 40059065-0) of University of O’Higgins; and Minera Valle Central company (Requinoa, O’Higgins, Chile).

## Supplementary information

The following supporting information can be downloaded:

– Draft_Systemix_MAGs_Profiling-Supplementary

## Supplementary tables

– Draft_Systemix_MAGs_Profiling-Supplementary.xlsx

## References

[1] S. Nayfach, Z. J. Shi, R. Seshadri, K. S. Pollard, and N. C. Kyrpides, “New insights from uncultivated genomes of the global human gut microbiome,” Nature, vol. 568, no. 7753, pp. 505–510, Apr. 2019, doi: 10.1038/s41586-019-1058-x.

[2] D. A. Cowan, J.-B. Ramond, T. P. Makhalanyane, and P. De Maayer, “Metagenomics of extreme environments,” Curr. Opin. Microbiol., vol. 25, pp. 97–102, Jun. 2015, doi: 10.1016/j.mib.2015.05.005.

[3] M. A. Sierra et al., “Microbiome and metagenomic analysis of Lake Hillier Australia reveals pigment-rich polyextremophiles and wide-ranging metabolic adaptations,” Environ Microbiome, vol. 17, no. 1, p. 60, Dec. 2022, doi: 10.1186/s40793-022-00455-9.

[4] V. Waschulin et al., “Biosynthetic potential of uncultured Antarctic soil bacteria revealed through long-read metagenomic sequencing,” ISME J., vol. 16, no. 1, pp. 101–111, Jan. 2022, doi: 10.1038/s41396-021-01052-3.

[5] S. Li et al., “Capturing the microbial dark matter in desert soils using culturomics-based metagenomics and high-resolution analysis,” NPJ Biofilms Microbiomes, vol. 9, no. 1, p. 67, Sep. 2023, doi: 10.1038/s41522-023-00439-8.

[6] J. M. Haro-Moreno, P. J. Cabello-Yeves, M. P. Garcillán-Barcia, A. Zakharenko, T. I. Zemskaya, and F. Rodriguez-Valera, “A novel and diverse group of Candidatus Patescibacteria from bathypelagic Lake Baikal revealed through long-read metagenomics,” Environ Microbiome, vol. 18, no. 1, p. 12, Feb. 2023, doi: 10.1186/s40793-023-00473-1.

[7] J.-L. Liang et al., “Hidden diversity and potential ecological function of phosphorus acquisition genes in widespread terrestrial bacteriophages,” Nat. Commun., vol. 15, no. 1, p. 2827, Apr. 2024, doi: 10.1038/s41467-024-47214-7.

[8] L. Zhang et al., “Bacterial and archaeal communities in the deep-sea sediments of inactive hydrothermal vents in the Southwest India Ridge,” Sci. Rep., vol. 6, p. 25982, May 2016, doi: 10.1038/srep25982.

[9] X. Dong et al., “Functional diversity of microbial communities in inactive seafloor sulfide deposits,” FEMS Microbiol. Ecol., vol. 97, no. 8, Aug. 2021, doi: 10.1093/femsec/fiab108.

[10] H. Zhao, Y. Li, M. Liu, X. Song, B. Liu, and X. Ju, “Metagenomic and network analyses reveal key players in nitrification in upland agricultural soils,” Environ. Microbiol., vol. 25, no. 11, pp. 2636–2640, Nov. 2023, doi: 10.1111/1462-2920.16467.

[11] M. Latorre et al., “The bioleaching potential of a bacterial consortium,” Bioresour. Technol., vol. 218, pp. 659–666, Oct. 2016, doi: 10.1016/j.biortech.2016.07.012.

[12] G. Galvez et al., “Co-occurrence Interaction Networks of Extremophile Species Living in a Copper Mining Tailing,” Front. Microbiol., vol. 12, p. 791127, 2021, doi: 10.3389/fmicb.2021.791127.

[13] J. Valdés et al., “Acidithiobacillus ferrooxidans metabolism: from genome sequence to industrial applications,” BMC Genomics, vol. 9, p. 597, Dec. 2008, doi: 10.1186/1471-2164-9-597.

[14] D. Travisany et al., “A new genome of Acidithiobacillus thiooxidans provides insights into adaptation to a bioleaching environment,” Res. Microbiol., vol. 165, no. 9, pp. 743–752, Nov. 2014, doi: 10.1016/j.resmic.2014.08.004.

[15] X. Zhang et al., “Comparative Genomics of the Extreme Acidophile Acidithiobacillus thiooxidans Reveals Intraspecific Divergence and Niche Adaptation,” Int. J. Mol. Sci., vol. 17, no. 8, Aug. 2016, doi: 10.3390/ijms17081355.

[16] R. Quatrini and D. B. Johnson, “Microbiomes in extremely acidic environments: functionalities and interactions that allow survival and growth of prokaryotes at low pH,” Curr. Opin. Microbiol., vol. 43, pp. 139–147, Jun. 2018, doi: 10.1016/j.mib.2018.01.011.

[17] P. R. Norris, D. A. Clark, J. P. Owen, and S. Waterhouse, “Characteristics of Sulfobacillus acidophilus sp. nov. and other moderately thermophilic mineral-sulphide-oxidizing bacteria,” Microbiology, vol. 142 ( Pt 4), pp. 775–783, Apr. 1996, doi: 10.1099/00221287-142-4-775.

[18] I. Anderson et al., “Complete genome sequence of the moderately thermophilic mineral-sulfide-oxidizing firmicute Sulfobacillus acidophilus type strain (NAL(T)),” Stand. Genomic Sci., vol. 6, no. 3, pp. 1–13, Jul. 2012, doi: 10.4056/sigs.2736042.

[19] L. Valenzuela et al., “Genomics, metagenomics and proteomics in biomining microorganisms,” Biotechnol. Adv., vol. 24, no. 2, pp. 197–211, Mar-Apr 2006, doi: 10.1016/j.biotechadv.2005.09.004.

[20] Y. Zhan et al., “Iron and sulfur oxidation pathways of Acidithiobacillus ferrooxidans,” World J. Microbiol. Biotechnol., vol. 35, no. 4, p. 60, Mar. 2019, doi: 10.1007/s11274-019-2632-y.

[21] M. Li and J. Wen, “Recent progress in the application of omics technologies in the study of bio-mining microorganisms from extreme environments,” Microb. Cell Fact., vol. 20, no. 1, p. 178, Sep. 2021, doi: 10.1186/s12934-021-01671-7.

[22] J. Gentina and F. Acevedo, “Copper Bioleaching in Chile,” Minerals (Basel), vol. 6, no. 1, p. 23, Mar. 2016, doi: 10.3390/min6010023.

[23] J. P. Cárdenas, R. Quatrini, and D. S. Holmes, “Genomic and metagenomic challenges and opportunities for bioleaching: a mini-review,” Res. Microbiol., vol. 167, no. 7, pp. 529–538, Sep. 2016, doi: 10.1016/j.resmic.2016.06.007.

[24] L. Yang et al., “Acidithiobacillus thiooxidans and its potential application,” Appl. Microbiol. Biotechnol., vol. 103, no. 19, pp. 7819–7833, Oct. 2019, doi: 10.1007/s00253-019-10098-5.

[25] P. Sharma, S. Kumar, and A. Pandey, “Bioremediated techniques for remediation of metal pollutants using metagenomics approaches: A review,” J. Environ. Chem. Eng., vol. 9, no. 4, p. 105684, Aug. 2021, doi: 10.1016/j.jece.2021.105684.

[26] W.-L. Huang, P.-C. Wu, and T.-Y. Chiang, “Metagenomics: Potential for bioremediation of soil contaminated with heavy metals,” Ecol. Genet. Genom., vol. 22, no. 100111, p. 100111, Mar. 2022, doi: 10.1016/j.egg.2021.100111.

[27] J. Chen, Y. Liu, P. Diep, and R. Mahadevan, “Genetic engineering of extremely acidophilic Acidithiobacillus species for biomining: Progress and perspectives,” J. Hazard. Mater., vol. 438, p. 129456, Sep. 2022, doi: 10.1016/j.jhazmat.2022.129456.

[28] R. Daniel, “The metagenomics of soil,” Nat. Rev. Microbiol., vol. 3, no. 6, pp. 470–478, Jun. 2005, doi: 10.1038/nrmicro1160.

[29] N. Fierer, “Embracing the unknown: disentangling the complexities of the soil microbiome,” Nat. Rev. Microbiol., vol. 15, no. 10, pp. 579–590, Oct. 2017, doi: 10.1038/nrmicro.2017.87.

[30] S. Banerjee and M. G. A. van der Heijden, “Soil microbiomes and one health,” Nat. Rev. Microbiol., vol. 21, no. 1, pp. 6–20, Jan. 2023, doi: 10.1038/s41579-022-00779-w.

[31] B. Ma et al., “A genomic catalogue of soil microbiomes boosts mining of biodiversity and genetic resources,” Nat. Commun., vol. 14, no. 1, p. 7318, Nov. 2023, doi: 10.1038/s41467-023-43000-z.

[32] D. C. Schlatter, K. Kahl, B. Carlson, D. R. Huggins, and T. Paulitz, “Soil acidification modifies soil depth-microbiome relationships in a no-till wheat cropping system,” Soil Biol. Biochem., vol. 149, no. 107939, p. 107939, Oct. 2020, doi: 10.1016/j.soilbio.2020.107939.

[33] J. Kumar, N. Sharma, and S. P. Singh, “Genome-resolved metagenomics inferred novel insights into the microbial community, metabolic pathways, and biomining potential of Malanjkhand acidic copper mine tailings,” Environ. Sci. Pollut. Res. Int., vol. 30, no. 17, pp. 50864–50882, Apr. 2023, doi: 10.1007/s11356-023-25893-x.

[34] S. Liu, C. D. Moon, N. Zheng, S. Huws, S. Zhao, and J. Wang, “Opportunities and challenges of using metagenomic data to bring uncultured microbes into cultivation,” Microbiome, vol. 10, no. 1, p. 76, May 2022, doi: 10.1186/s40168-022-01272-5.

[35] L.-X. Chen et al., “Comparative metagenomic and metatranscriptomic analyses of microbial communities in acid mine drainage,” ISME J., vol. 9, no. 7, pp. 1579–1592, Jul. 2015, doi: 10.1038/ismej.2014.245.

[36] S. Krakau, D. Straub, H. Gourlé, G. Gabernet, and S. Nahnsen, “nf-core/mag: a best-practice pipeline for metagenome hybrid assembly and binning,” NAR Genom Bioinform, vol. 4, no. 1, p. lqac007, Mar. 2022, doi: 10.1093/nargab/lqac007.

[37] P. Di Tommaso, M. Chatzou, E. W. Floden, P. P. Barja, E. Palumbo, and C. Notredame, “Nextflow enables reproducible computational workflows,” Nat. Biotechnol., vol. 35, no. 4, pp. 316–319, Apr. 2017, doi: 10.1038/nbt.3820.

[38] R. M. Bowers et al., “Minimum information about a single amplified genome (MISAG) and a metagenome-assembled genome (MIMAG) of bacteria and archaea,” Nat. Biotechnol., vol. 35, no. 8, pp. 725–731, Aug. 2017, doi: 10.1038/nbt.3893.

[39] D. H. Parks et al., “A standardized bacterial taxonomy based on genome phylogeny substantially revises the tree of life,” Nat. Biotechnol., vol. 36, no. 10, pp. 996–1004, Nov. 2018, doi: 10.1038/nbt.4229.

[40] P.-A. Chaumeil, A. J. Mussig, P. Hugenholtz, and D. H. Parks, “GTDB-Tk: a toolkit to classify genomes with the Genome Taxonomy Database,” Bioinformatics, vol. 36, no. 6, pp. 1925–1927, Nov. 2019, doi: 10.1093/bioinformatics/btz848.

[41] T. D’Angelo et al., “Replicated life-history patterns and subsurface origins of the bacterial sister phyla Nitrospirota and Nitrospinota,” ISME J., vol. 17, no. 6, pp. 891–902, Jun. 2023, doi: 10.1038/s41396-023-01397-x.

[42] K. Umezawa, H. Kojima, Y. Kato, and M. Fukui, “Disproportionation of inorganic sulfur compounds by a novel autotrophic bacterium belonging to Nitrospirota,” Syst. Appl. Microbiol., vol. 43, no. 5, p. 126110, Sep. 2020, doi: 10.1016/j.syapm.2020.126110.

[43] N. Kochhar et al., “Perspectives on the microorganism of extreme environments and their applications,” Curr Res Microb Sci, vol. 3, p. 100134, Apr. 2022, doi: 10.1016/j.crmicr.2022.100134.

[44] M.-G. Sánchez-Otero, R. Quintana-Castro, J. G. Domínguez-Chávez, C. Peña-Montes, and R. M. Oliart-Ros, “Unique microorganisms inhabit extreme soils,” in Microorganisms for Sustainability, Singapore: Springer Singapore, 2019, pp. 39–73. doi: 10.1007/978-981-13-9117-0_3.

[45] M. Delgado-Baquerizo et al., “A global atlas of the dominant bacteria found in soil,” Science, vol. 359, no. 6373, pp. 320–325, Jan. 2018, doi: 10.1126/science.aap9516.

[46] X. Li et al., “Acidification suppresses the natural capacity of soil microbiome to fight pathogenic Fusarium infections,” Nat. Commun., vol. 14, no. 1, p. 5090, Aug. 2023, doi: 10.1038/s41467-023-40810-z.

[47] J.-T. Li et al., “Metagenomic and metatranscriptomic insights into sulfate-reducing bacteria in a revegetated acidic mine wasteland,” NPJ Biofilms Microbiomes, vol. 8, no. 1, p. 71, Sep. 2022, doi: 10.1038/s41522-022-00333-9.

[48] L. P. M. Lamers et al., “Microbial transformations of nitrogen, sulfur, and iron dictate vegetation composition in wetlands: a review,” Front. Microbiol., vol. 3, p. 156, Apr. 2012, doi: 10.3389/fmicb.2012.00156.

[49] M. Miransari, “Soil microbes and the availability of soil nutrients,” Acta Physiol. Plant, vol. 35, no. 11, pp. 3075–3084, Nov. 2013, doi: 10.1007/s11738-013-1338-2.

[50] G. Feng et al., “Metagenomic analysis of microbial community and function involved in cd-contaminated soil,” BMC Microbiol., vol. 18, no. 1, p. 11, Feb. 2018, doi: 10.1186/s12866-018-1152-5.

[51] V. De Anda, I. Zapata-Peñasco, A. C. Poot-Hernandez, L. E. Eguiarte, B. Contreras-Moreira, and V. Souza, “MEBS, a software platform to evaluate large (meta)genomic collections according to their metabolic machinery: unraveling the sulfur cycle,” Gigascience, vol. 6, no. 11, pp. 1–17, Nov. 2017, doi: 10.1093/gigascience/gix096.

[52] J. Mistry et al., “Pfam: The protein families database in 2021,” Nucleic Acids Res., vol. 49, no. D1, pp. D412–D419, Jan. 2021, doi: 10.1093/nar/gkaa913.

[53] A. Levett, S. A. Gleeson, and J. Kallmeyer, “From exploration to remediation: A microbial perspective for innovation in mining,” Earth Sci. Rev., vol. 216, no. 103563, p. 103563, May 2021, doi: 10.1016/j.earscirev.2021.103563.

[54] I. R. Ugwuanyi et al., “Comparative metagenomics at Solfatara and Pisciarelli hydrothermal systems in Italy reveal that ecological differences across substrates are not ubiquitous,” Front. Microbiol., vol. 14, p. 1066406, Feb. 2023, doi: 10.3389/fmicb.2023.1066406.

[55] D. J. Levy-Booth, C. E. Prescott, and S. J. Grayston, “Microbial functional genes involved in nitrogen fixation, nitrification and denitrification in forest ecosystems,” Soil Biol. Biochem., vol. 75, pp. 11–25, Aug. 2014, doi: 10.1016/j.soilbio.2014.03.021.

[56] S. Pandey, E. Fosso-Kankeu, J. Redelinghuys, J. Kim, and M. Kang, “Implication of biofilms in the sustainability of acid mine drainage and metal dispersion near coal tailings,” Sci. Total Environ., vol. 788, p. 147851, Sep. 2021, doi: 10.1016/j.scitotenv.2021.147851.

[57] P. Vidal et al., “Metagenomic Mining for Esterases in the Microbial Community of Los Rueldos Acid Mine Drainage Formation,” Front. Microbiol., vol. 13, p. 868839, May 2022, doi: 10.3389/fmicb.2022.868839.

[58] Z. She et al., “Multi-omics insights into biogeochemical responses to organic matter addition in an acidic pit lake: Implications for bioremediation,” Water Res., vol. 254, p. 121404, May 2024, doi: 10.1016/j.watres.2024.121404.

[59] D. Liao, “Gene conversion drives within genic sequences: concerted evolution of ribosomal RNA genes in bacteria and archaea,” J. Mol. Evol., vol. 51, no. 4, pp. 305–317, Oct. 2000, doi: 10.1007/s002390010093.

[60] W. K. Ray, G. Zeng, M. B. Potters, A. M. Mansuri, and T. J. Larson, “Characterization of a 12-kilodalton rhodanese encoded by glpE of Escherichia coli and its interaction with thioredoxin,” J. Bacteriol., vol. 182, no. 8, pp. 2277–2284, Apr. 2000, doi: 10.1128/JB.182.8.2277-2284.2000.

[61] M. Lu, J. Chai, and D. Fu, “Structural basis for autoregulation of the zinc transporter YiiP,” Nat. Struct. Mol. Biol., vol. 16, no. 10, pp. 1063–1067, Oct. 2009, doi: 10.1038/nsmb.1662.

[62] J. Sun et al., “Comparative genomics provides new insights into pathogenesis of spotting disease causative bacteria in farmed Strongylocentrotus intermedius,” Aquaculture, vol. 539, no. 736564, p. 736564, Jun. 2021, doi: 10.1016/j.aquaculture.2021.736564.

[63] J. J. Harrison, H. Ceri, and R. J. Turner, “Multimetal resistance and tolerance in microbial biofilms,” Nat. Rev. Microbiol., vol. 5, no. 12, pp. 928–938, Dec. 2007, doi: 10.1038/nrmicro1774.

[64] S.-J. Li et al., “Microbial communities evolve faster in extreme environments,” Sci. Rep., vol. 4, p. 6205, Aug. 2014, doi: 10.1038/srep06205.

[65] S. Turner, K. M. Pryer, V. P. Miao, and J. D. Palmer, “Investigating deep phylogenetic relationships among cyanobacteria and plastids by small subunit rRNA sequence analysis,” J. Eukaryot. Microbiol., vol. 46, no. 4, pp. 327–338, Jul-Aug 1999, doi: 10.1111/j.1550-7408.1999.tb04612.x.

[66] S. Chen, Y. Zhou, Y. Chen, and J. Gu, “fastp: an ultra-fast all-in-one FASTQ preprocessor,” Bioinformatics, vol. 34, no. 17, pp. i884–i890, Sep. 2018, doi: 10.1093/bioinformatics/bty560.

[67] A. Bankevich et al., “SPAdes: a new genome assembly algorithm and its applications to single-cell sequencing,” J. Comput. Biol., vol. 19, no. 5, pp. 455–477, May 2012, doi: 10.1089/cmb.2012.0021.

[68] A. Prjibelski, D. Antipov, D. Meleshko, A. Lapidus, and A. Korobeynikov, “Using SPAdes De Novo Assembler,” Curr. Protoc. Bioinformatics, vol. 70, no. 1, p. e102, Jun. 2020, doi: 10.1002/cpbi.102.

[69] D. D. Kang et al., “MetaBAT 2: an adaptive binning algorithm for robust and efficient genome reconstruction from metagenome assemblies,” PeerJ, vol. 7, p. e7359, Jul. 2019, doi: 10.7717/peerj.7359.

[70] Y.-W. Wu, B. A. Simmons, and S. W. Singer, “MaxBin 2.0: an automated binning algorithm to recover genomes from multiple metagenomic datasets,” Bioinformatics, vol. 32, no. 4, pp. 605–607, Feb. 2016, doi: 10.1093/bioinformatics/btv638.

[71] C. M. K. Sieber et al., “Recovery of genomes from metagenomes via a dereplication, aggregation and scoring strategy,” Nat Microbiol, vol. 3, no. 7, pp. 836–843, Jul. 2018, doi: 10.1038/s41564-018-0171-1.

[72] D. Albanese and C. Donati, “Large-scale quality assessment of prokaryotic genomes with metashot/prok-quality,” F1000Res., vol. 10, p. 822, Aug. 2021, doi: 10.12688/f1000research.54418.1.

[73] D. H. Parks, M. Imelfort, C. T. Skennerton, P. Hugenholtz, and G. W. Tyson, “CheckM: assessing the quality of microbial genomes recovered from isolates, single cells, and metagenomes,” Genome Res., vol. 25, no. 7, pp. 1043–1055, Jul. 2015, doi: 10.1101/gr.186072.114.

[74] D. Hyatt, G.-L. Chen, P. F. Locascio, M. L. Land, F. W. Larimer, and L. J. Hauser, “Prodigal: prokaryotic gene recognition and translation initiation site identification,” BMC Bioinformatics, vol. 11, p. 119, Mar. 2010, doi: 10.1186/1471-2105-11-119.

[75] O. Schwengers, L. Jelonek, M. A. Dieckmann, S. Beyvers, J. Blom, and A. Goesmann, “Bakta: rapid and standardized annotation of bacterial genomes via alignment-free sequence identification,” Microb Genom, vol. 7, no. 11, Nov. 2021, doi: 10.1099/mgen.0.000685.

[76] G. Yu, D. K. Smith, H. Zhu, Y. Guan, and T. T.-Y. Lam, “Ggtree: An r package for visualization and annotation of phylogenetic trees with their covariates and other associated data,” Methods Ecol. Evol., vol. 8, no. 1, pp. 28–36, Jan. 2017, doi: 10.1111/2041-210x.12628.

[77] T. Aramaki et al., “KofamKOALA: KEGG Ortholog assignment based on profile HMM and adaptive score threshold,” Bioinformatics, vol. 36, no. 7, pp. 2251–2252, Apr. 2020, doi: 10.1093/bioinformatics/btz859.

[78] Z. Gu, R. Eils, and M. Schlesner, “Complex heatmaps reveal patterns and correlations in multidimensional genomic data,” Bioinformatics, vol. 32, no. 18, pp. 2847–2849, Sep. 2016, doi: 10.1093/bioinformatics/btw313.

[79] M. Vasimuddin, S. Misra, H. Li, and S. Aluru, “Efficient architecture-aware acceleration of BWA-MEM for multicore systems,” in 2019 IEEE International Parallel and Distributed Processing Symposium (IPDPS), IEEE, May 2019. doi: 10.1109/ipdps.2019.00041.

[80] H. Li et al., “The Sequence Alignment/Map format and SAMtools,” Bioinformatics, vol. 25, no. 16, pp. 2078–2079, Aug. 2009, doi: 10.1093/bioinformatics/btp352.

[81] C. Herzeel, P. Costanza, D. Decap, J. Fostier, R. Wuyts, and W. Verachtert, “Multithreaded variant calling in elPrep 5,” PLoS One, vol. 16, no. 2, p. e0244471, Feb. 2021, doi: 10.1371/journal.pone.0244471.

[82] P. Danecek et al., “Twelve years of SAMtools and BCFtools,” Gigascience, vol. 10, no. 2, Feb. 2021, doi: 10.1093/gigascience/giab008.

[83] P. Cingolani et al., “A program for annotating and predicting the effects of single nucleotide polymorphisms, SnpEff: SNPs in the genome of Drosophila melanogaster strain w1118; iso-2; iso-3,” Fly, vol. 6, no. 2, pp. 80–92, Apr-Jun 2012, doi: 10.4161/fly.19695.

[84] I. Martincorena et al., “Universal Patterns of Selection in Cancer and Somatic Tissues,” Cell, vol. 171, no. 5, pp. 1029–1041.e21, Nov. 2017, doi: 10.1016/j.cell.2017.09.042.

[85] Z. Gu, L. Gu, R. Eils, M. Schlesner, and B. Brors, “circlize Implements and enhances circular visualization in R,” Bioinformatics, vol. 30, no. 19, pp. 2811–2812, Oct. 2014, doi: 10.1093/bioinformatics/btu393.

