## Supplementary Material for "Profiling extremophile bacterial communities recovered from a mining tailing against soil ecosystems through comparative genome-resolved metagenomics and evolutionary analysis"

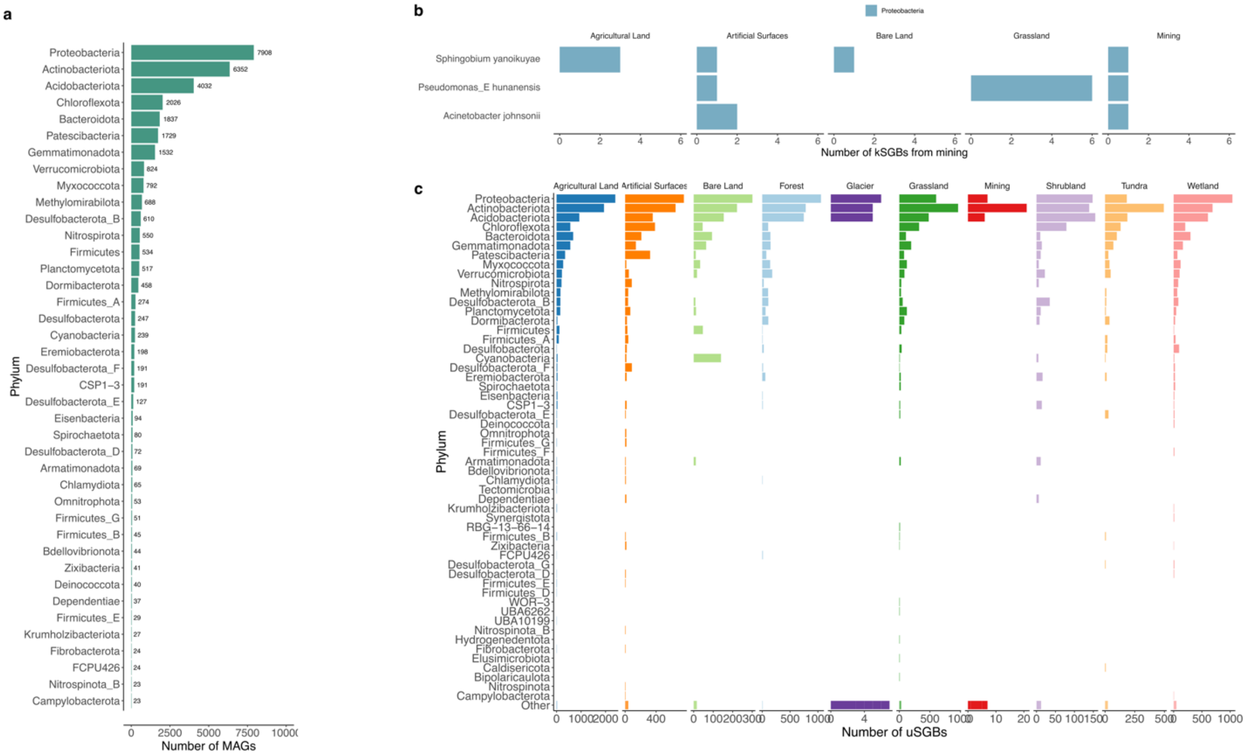


**Supplementary Figure 1. a)** Ranked histogram of the top 40 most abundant phylum in the combined mining MAGs and the SMAG catalog. **b)** Number of known species (kSGBs) from the mining ecosystem and the conventional ecosystems in which they were also found. **c)** Distribution of unknown species (uSGBs) across mining and conventional ecosystems. The phyla with low counts are summarized in the last row as “other”.

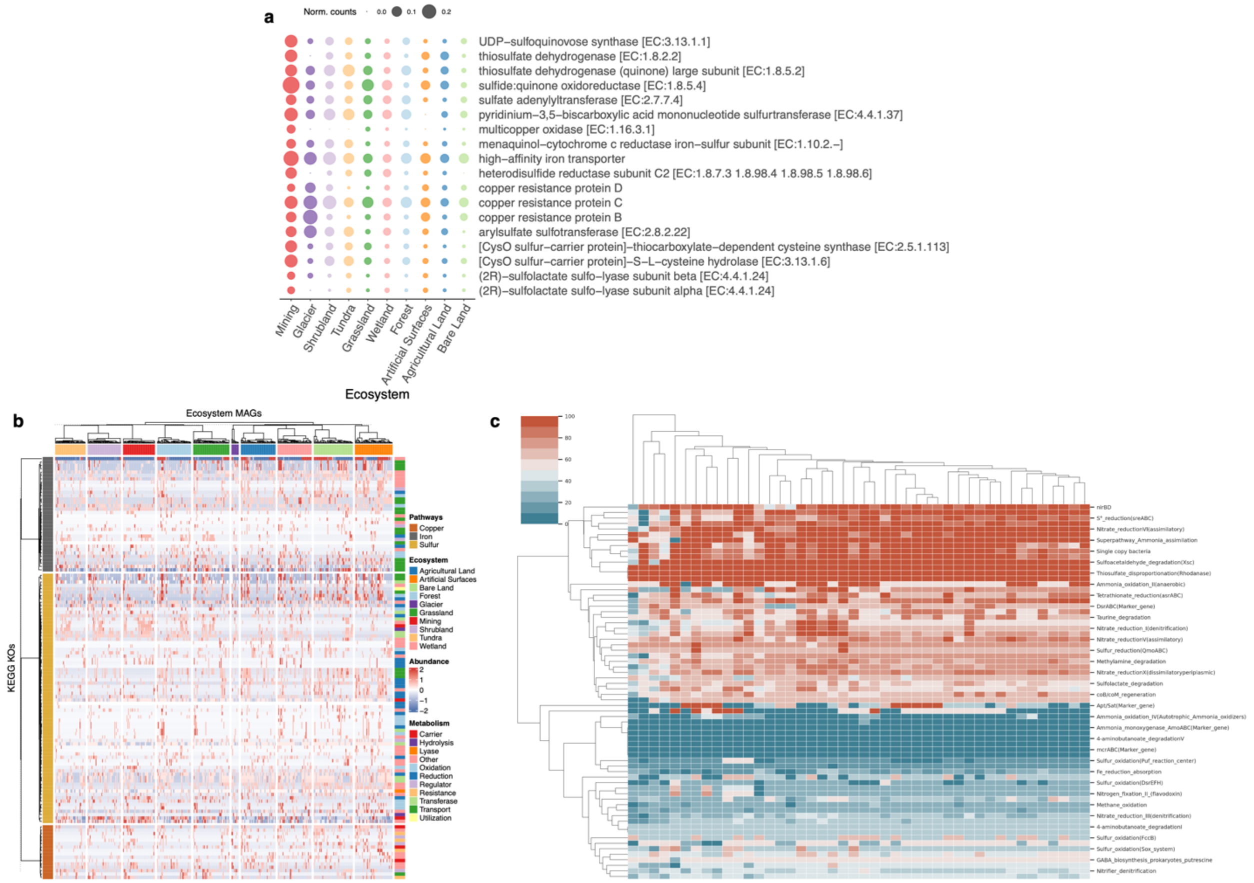


**Supplementary Figure 2. a)** Bubble plot of most abundant genes from mining MAGs, where the bigger circle of normalized counts denote higher information of each KO definition in their respective ecosystem. **b)** Annotated heatmap of KEGG KOs across all ecosystems (in columns), labeling the selected pathways (copper, iron and sulfur in left rows annotation) and their main metabolism (left rows annotation). Values are normalized by z-score and the red color indicates high prevalence of a given MAG in the target pathway (abundances). Dendrograms in rows and columns reflect similarity among KOs and ecosystems, respectively. **c)** Heatmap of mining MAGs metabolism annotated with MEBS. Red color indicates high likelihood of a given genome at that pathway, meaning higher information in the target metabolism (y-axis).

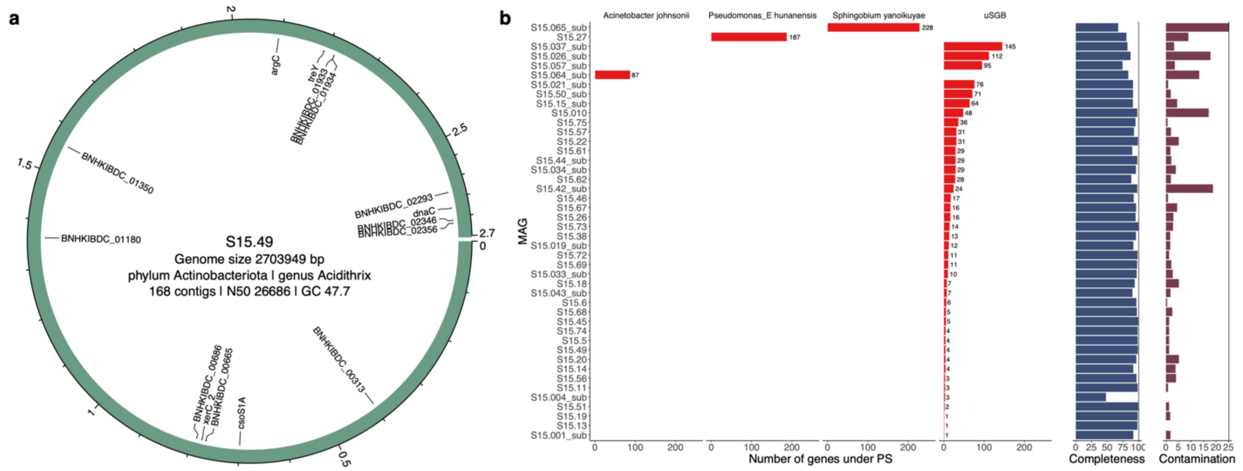


**Supplementary Figure 3. a)** Circos plot representation of the mining MAG with global positive selection depicting the position of annotated genes under PS. **b)** Summary of genes under PS grouped and sorted by MAG at species-level of classification, including both completeness and contamination from QC. Each bin contains at least 1 SNP.

**Supplementary Table 1.** Global statistics of the sequenced metagenomic sample, parameters after reads filtering and their assembly.

| Sample | Raw reads | Length (Gbp) | Passed filters (%) | Duplication (%) | GC (%) | # contigs | N50 (kbp) |
|---|---|---|---|---|---|---|---|
| S15 | 103,790,393 | 31.13 | 99.2 | 14.9 | 59.1 | 573,859 | 9.0 |
